# iTaxoTools 0.1: Kickstarting a specimen-based software toolkit for taxonomists

**DOI:** 10.1101/2021.03.26.435825

**Authors:** Miguel Vences, Aurélien Miralles, Sophie Brouillet, Jacques Ducasse, Alexander Fedosov, Vladimir Kharchev, Ivaylo Kostadinov, Sangeeta Kumari, Stefanos Patmanidis, Mark D. Scherz, Nicolas Puillandre, Susanne S. Renner

## Abstract

While powerful and user-friendly software suites exist for phylogenetics, and an impressive cybertaxomic infrastructure of online species databases has been set up in the past two decades, software specifically targeted at facilitating alpha-taxonomic work, i.e., delimiting and diagnosing species, is still in its infancy. Here we present a project to develop a bioinformatic toolkit for taxonomy, based on open-source Python code, including tools focusing on species delimitation and diagnosis and centered around specimen identifiers. At the core of iTaxoTools is user-friendliness, with numerous autocorrect options for data files and with intuitive graphical user interfaces. Assembled standalone executables for all tools or a suite of tools with a launcher window will be distributed for Windows, Linux, and Mac OS systems, and in the future also implemented on a web server. The alpha version (iTaxoTools 0.1) distributed with this paper contains GUI versions of six species delimitation programs (ABGD, ASAP, DELINEATE, GMYC, PTP, tr2) and a simple threshold-clustering delimitation tool. There are also new Python implementations of existing algorithms, including tools to compute pairwise DNA distances, ultrametric time trees based on non-parametric rate smoothing, species-diagnostic nucleotide positions, and standard morphometric analyses. Other utilities convert among different formats of molecular sequences, geographical coordinates, and units; merge, split and prune sequence files and tables; and perform simple statistical tests. As a future perspective, we envisage iTaxoTools to become part of a bioinformatic pipeline for next-generation taxonomy that accelerates the inventory of life while maintaining high-quality species hypotheses.

## Introduction

Bioinformatics has become the core of modern biology, especially in the context of high-throughput workflows that are becoming commonplace in many fields, in particular related to - omics approaches. The big data volumes obtained by these techniques require ever more efficient and sophisticated software, which is being developed and refined at a vigorous pace. In the field of systematics, powerful programs for phylogenetic analysis have been developed, and databases and data aggregators have been set up to deal with the massive globally-generated taxonomic dataset comprised of over one million species and many millions of specimen records. Yet, only few bioinformatic tools so far have been tailored to specifically fit the practical work of taxonomists, who diagnose and name some 15,000–20,000 new species of organisms per year, a task that still is largely performed by single or small teams of (professional and amateur) researchers (Miralles et al. 2020). Most existing tools are aimed at the construction of identification keys (e.g. Dallwitz, 1974; Clark 2003; Delgado Calvo-Flores et al. 2006; Zhang et al. 2006; MacLeod 2008; Vignes Lebbe 2015; Tofilski 2018), which in some groups help field identification. Only a handful of software packages (EDIT: cybertaxonomy.eu, TaxonWorks: taxonworks.org, Scratchpads: scratchpads.org) are tailored towards facilitating descriptive work itself, but none of these is so far widely used; furthermore, these programs do not include various important aspects of the alpha-taxonomic workflow, such as species delimitation or molecular diagnosis (Miralles et al. 2020) which can also be of high relevance for other fields such as molecular ecology.

Although most taxonomic studies are still relying on morphology only (as shown in a recent review; Miralles et al. 2020), taxonomy increasingly integrates diverse lines of evidence (Padial et al. 2010), a procedure called integrative taxonomy by Dayrat (2005). Discovering, delimiting, diagnosing, and naming new species requires taxonomists to examine voucher specimens and associated catalogues, field books and pictures; take, tabulate and statistically analyze morphometric measurements; define, tabulate and document phenotypic character states; estimate geographical ranges based on specimen provenances; align and analyze DNA sequences; and elaborate accurate specimen tables, species diagnoses and identification keys. Depending on the organism under study, it also may involve more specialized procedures such as comparing acoustic and visual signal repertoires of animals, or isolate and culture unicellular organisms. In addition, to fulfil standards of cybertaxonomy, data sets need to be archived in specialized repositories and new species names registered in online databases (Miralles et al. 2020). With rising best-practice standards, these many and varied tasks generally involve the use of different computer programs – and thus lead to an extra burden on taxonomists who may lack bioinformatic training.

## The concept of iTaxoTools

We aim to develop a bioinformatic platform to facilitate the core work of taxonomists, that is, delimiting, diagnosing and describing species. Our initiative produced an integrative taxonomy toolkit – iTaxoTools (Fig. 1; Table 1). The concept of iTaxoTools rests on four pillars: (1) **fully open source** code; (2) a **diversified** set of stand-alone programs (‘modules’) that in future versions will become increasingly interconnected; (3) a **specimen-centered** architecture, where at present tables (tab-delimited text files) with specimen identifier columns serve as main input format; and (4) a focus on **user-friendliness**, accessibility, and clear and transparent documentation.

**FIGURE 1.**
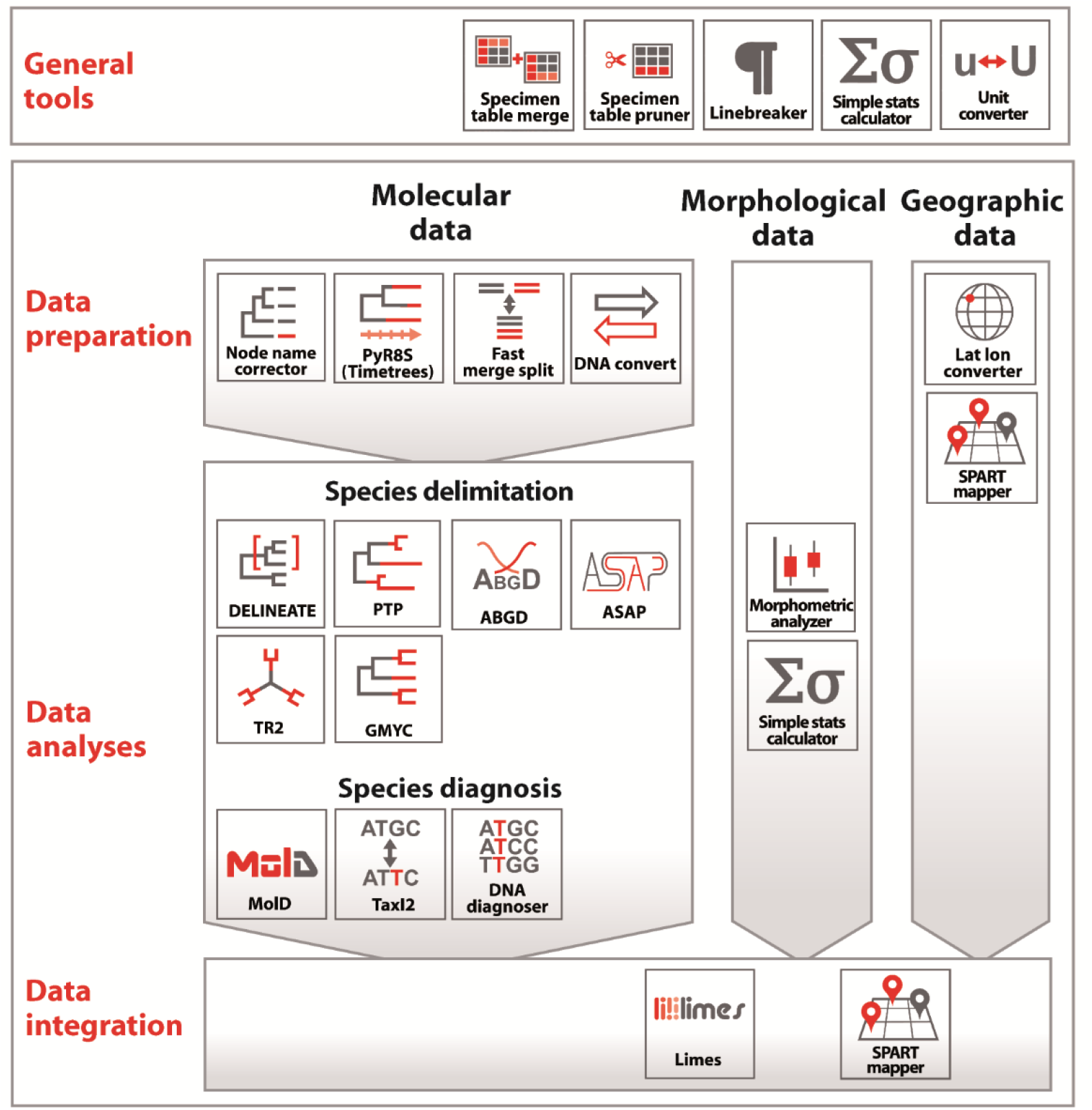
Overview of the various tools implemented in iTaxoTools, and their scope. In the present version a focus is on molecular data analysis, but more functionalities to analyze and visualize morphological and geographic data will be implemented in the next future, while data integration remains the main focus for long-term implementation.

**TABLE 1.** Overview of the software tools and functionalities currently included in the 0.1 version of iTaxoTools. The majority of the tools can be run (i) command-line driven in Python, and is distributed as (ii) a standalone executable (.exe) file, (iii) as part of a full package with launcher window (Fig. 1) in a single executable or part of a folder, and (iv) all tools will furthermore be implemented on a webserver. Note that functionalities for pre-existing species delimitation tools are explained in more detail in the original papers.

| Tool | Purpose | Main functionalities |
| --- | --- | --- |
| dnaconvert | Converts among DNA sequence formats | - Supports typical sequence formats (fasta, fastq, phylip, nexus)<br>- Autocorrects typical errors in sequence files such as non-standard characters in sequence names.<br>- Reads GenBank flat files and converts from and to tab-delimited files to manage sequence in spreadsheet editors.<br>- Single-file conversion, batch conversion and conversion of copy-pasted files |
| latlonconverter | Converts among different geographic coordinate formats | - Parses a large variety of formats of WGS84 geographical coordinates<br>- Batch-conversion of coordinates in tables or copy-pasted lists which can contain coordinates in different formats (recognized by heuristic approach)<br>- Main output in decimal degree format |
| fastmerge | Merges DNA sequence files (fasta, fastq) | - Can merge large /definition?/ files that usually cannot be opened in editors.<br>- Works for any text file but includes additional features when processing fasta and fastq.<br>- Allows for filtering sequences and sequence names with certain motifs and include/exclude them in the merged file |
| fastsplit | Splits (large) DNA sequence files (fasta, fastq) into smaller files | - Can split large files /definition?/ that usually cannot be opened in editors and split them into a series of equally sized smaller files<br>- Designed for fasta and fastq, but works for any text file.<br>- Allows for filtering sequences and sequence names with certain motifs and include/exclude them in the split files |
| specimentablepruner | Removes rows from tables based on a list of values for the row "specimen" | - Takes as input a tab-delimited file and a series of values of specimen identifiers<br>- Removes all rows from the table where the column "specimen" (or other chosen column) agrees with any of the provided values |
| specimentablemerger | Merges data from two tables based on values in the row "specimen" | - Takes as input two or more tab-delimited files, compares values in column "specimen" (or other chosen column) and merges into one table, combining values for same specimen number in the same row<br>- Automatically checks for duplicate values of the same variable and specimen and issues warnings |
| linebreaker | Changes among line break formats (Unix, Windows, Mac) | - Takes as input any text file and changes all line breaks to the specified format |
| simplestatscalculator | Performs a series of basic statistical analyses based manually entered data | - Values are typed or pasted into text boxes<br>- Descriptive statistics (mean, median, standard deviation and others)<br>- Pairwise comparisons (t-test, U-test)<br>- Comparisons of distributions (Chi-square, normality, Fisher's)<br>- Corrections for multiple testing |
| unitconverter | Converts among different units | - Values are typed into one field, all other fields show converted values in real time<br>- Separate tabs for conversion of distance, volume, time, molarity, and others |
| spartmapper | Computes a kml file from geographical coordinates and spart file | - Takes as input a text file with decimal geographical coordinates and specimen identifiers, and a species partition (spart) file<br>- Outputs a kml file to show localities by species on Google Earth |
| nodenamecorrector | Removes special characters from terminal node names in Newick-formatted trees | - Takes as input a Newick treefile, identifies node names, searches for characters in node names that are not standard alphanumerical, and replaces them with underscores |
| pyr8s | Calculates ultrametric timetrees | - Takes as input treefiles with branch length (phylograms) |
|  | (chronograms) based on non-parametric rate-smoothing | - Transforms into ultrametric using non-parametric rate smoothing, without the need to access original data (sequences)<br>- User-friendly interface to set time constraints (calibrations) on nodes. |
| TaxI2 | Calculates inter- and intraspecific distances and the barcoding gap based on pairwise-aligning DNA sequences | - Takes as input aligned or unaligned sequence files in fasta or tab-delimited text format<br>- For unaligned sequences, pairwise alignment are performed<br>- Calculates pairwise genetic distances among all sequences<br>- If tab file contains row with species names, inter- and intra-species distances are calculated and summarized, and the barcoding gap as well as some summary statistics of the barcoding gap calculated<br>- inter-species distances are separately calculated for species of the same genus vs. different genera<br>- A histogram with illustrating the barcoding gap is produced in editable PDF format |
| morphometricanalyzer | Calculates a series of basic statistical comparisons of species based on morphometric data | - Takes as input tab-delimited files with morphometric measurements (continuous variables)<br>- Allows specifying if analyses should be done by species, by sex/stage, or by species and sex/stage<br>- Calculates summary statistics, pairwise comparisons (t-tests, U-tests), ANOVAs, PCA and DA<br>- Size-corrects values by calculating ratio against a reference measurement such as body size<br>- Outputs boxplots and scatterplots of PCA and DA factors, by species and/or sex/stage in editable PDF format<br>- Writes text output summarizing diagnostic characters (scientifically different measurements between species, with and without overlap of ranges) |
| dnadiagnoser | Computes diagnostic sites for species from DNA sequences | - Takes as input aligned or unaligned sequence files in fasta or tab-delimited text format<br>- Unaligned sequences are pairwise aligned to reference sequence and differences recorded relative to position in reference<br>- Summarizes variation within species and outputs diagnostic sites among species<br>- Outputs unique diagnostic sites for the whole data sets, as well as diagnostic sites in pairwise comparisons among species<br>- Output is given in the form of tables but also as text which can be used for species diagnoses in taxonomic papers |
| PTP | Species delimitation based on Poisson tree processes | - Uses as input a non-ultrametric tree with branch lengths (phylogram) tree in Newick or Nexus format<br>- Models speciation on branching events in terms of number of mutations (inferred from branch lengths)<br>- Bayesian and ML versions of PTP are implemented |
| GMYC | Species delimitation based on the Generalized Mixed Yule Coalescent | - Uses as input an ultrametric tree in Newick or Nexus format<br>- Uses a likelihood approach to analyse the timing of branching events, seeking for significant switches between a Yule (interspecific) and a coalescent (intraspecific) branching structure. |
| tr2 | Species delimitation using Bayesian model comparison and rooted triplets | - Takes as input a set of gene trees, and optionally a guide species tree<br>- Calculates posterior probability scores for user-specified delimitation hypotheses.<br>- Alternatively, finds the best delimitation under a guide tree specifying a hierarchical structure of species grouping. |
| DELINEATE | Species delimitation by integrating an explicit model of speciation into the multipopulation coalescent | - Takes as input a rooted ultrametric tree from a multispecies coalescent analysis, in Nexus or Newick format<br>- Second input file is a table assigning specimens to species, or flagging their species identity as unknown<br>- Outputs various alternative species partitions, ranked by probability |
| ABGD | Species delimitation by automatic barcoding gap discovery | - Takes as input a set of aligned sequences and calculates pairwise distances<br>- Uses a coalescent model to identify the position of the most likely barcode gap, based on a maximal genetic intraspecific divergence defined a priori by the user.<br>- Uses the DNA barcoding gap to propose species partitions. |
| ASAP | Species delimitation from single-locus sequence data by the Assemble Species by Automatic Partitioning approach | - Takes as input a set of aligned sequences and calculates pairwise distances<br>- Proposes species partitions ranked by a new scoring system that uses no biological prior insight of intraspecific diversity. |
| LIMES 2.0 | Compare species partitions by different indexes and parsing/merge/export spart files | - Reads species partition (spart) files, as well as species partition information in spreadsheet format<br>- Computes $C_{tax}$, $mC_{tax}$, $R_{tax}$ and Match Ratio indexes<br>- Can merge, extract and export spart files |
| MolD | Recovers DNA-based diagnoses for taxa from DNA sequence alignments | - recovers diagnostic combinations of nucleotides (DNCs) for pre-defined groups of DNA sequences, corresponding to taxa<br>- Identifies pure diagnostic sites, minimal DNCs (mDNCs), and redundant DNCs (rDNCs), the latter fulfil predefined criteria of reliability |

All of the code developed by us is **fully open source** and available from a dedicated GitHub repository (https://github.com/iTaxoTools). In the case of tools programmed by other researchers, we make this information transparent, and the GUI we added specifies the original references and programmers. The current pre-release of compiled executables is available from https://github.com/iTaxoTools/iTaxoTools-Executables. See Table 2 for repositories of each single tool.

**TABLE 2.** Repositories of the code of the tools included in the 0.1 version of iTaxoTools. The table also lists the main programmers involved in the development of each tool or its graphical user interface (GUI), and informs whether a tool was newly programmed for this project, adjusted from existing code (by adding a GUI plus sometimes additional functionalities), or included as original code and GUI without modification.

| Tool | New / Adjusted / Original | Github repository (original / modified) | Main programmers (original program) / GUI |
| --- | --- | --- | --- |
| dnaconvert | New | https://github.com/iTaxoTools/DNAconvert | V. Kharchev |
| latlonconverter | New | https://github.com/iTaxoTools/latlon-converter | V. Kharchev |
| fastmerge | New | https://github.com/iTaxoTools/fastsplit-merge | V. Kharchev |
| fastsplit | New | https://github.com/iTaxoTools/fastsplit-merge | V. Kharchev |
| specimentablepruner | New | https://github.com/iTaxoTools/specimentablepruner | V. Kharchev |
| specimentablemerger | New | https://github.com/iTaxoTools/specimentablemerger | V. Kharchev |
| linebreaker | New | https://github.com/iTaxoTools/linebreaker | S. Kumari |
| simplestatscalculator | New | https://github.com/iTaxoTools/simple_stat | S. Kumari |
| unitconverter | New | https://github.com/iTaxoTools/UnitConverter | S. Kumari |
| spartmapper | New | https://github.com/iTaxoTools/linebreak_replacer | S. Kumari |
| nodenamecorrector | New | https://github.com/iTaxoTools/nodenamecorrector | V. Kharchev |
| pyr8s | New | https://github.com/iTaxoTools/pyr8s | S. Patmanidis |
| TaxI2 | New | https://github.com/iTaxoTools/TaxI2 | V. Kharchev |
| morphometricanalyzer | New | https://github.com/iTaxoTools/morphometricanalyzer | V. Kharchev |
| dnadiagnoser | New | https://github.com/iTaxoTools/dnadiagnoser | V. Kharchev |
| PTP | Adjusted | https://github.com/zhangjiajie/PTP<br>https://github.com/iTaxoTools/PTP-PYQT5 | (J. Zhang)<br>GUI: S. Kumari |
| GMYC | Adjusted | https://github.com/zhangjiajie/pGMYC<br>https://github.com/iTaxoTools/GMYC-PYQT5 | (J. Zhang )<br>GUI: S. Kumari |
| tr2 | Adjusted | https://github.com/tfujisawa/tr2-delimitation-git<br>https://github.com/iTaxoTools/pyqt5-tr2 | (T. Fujisawa)<br>GUI: S. Kumari |
| DELINEATE | Adjusted | https://github.com/iTaxoTools/pyqt5-delineate | (J. Sukumaran)<br>GUI: S. Kumari |
| ABGD | Adjusted | https://bioinfo.mnhn.fr/abi/public/abgd/<br>https://github.com/iTaxoTools/ABGDpy | (S. Brouillet)<br>GUI: S. Patmanidis |
| ASAP | Adjusted | https://bioinfo.mnhn.fr/abi/public/asap/<br>https://github.com/iTaxoTools/ASAPy | (S. Brouillet)<br>GUI: S. Patmanidis |
| LIMES 2.0 | Original | https://github.com/iTaxoTools/LIMES | J. Ducasse |
| MolD | Adjusted | https://github.com/SashaFedosov/MolD<br>https://github.com/iTaxoTools/MolD_pyqt5 | (A. Fedosov)<br>GUI: S. Kumari |

The toolkit is **diversified**, including simple format converters of molecular or geographic data, text and spreadsheet merging and pruning, simple statistical analyses e.g. of morphometric data, but especially focuses on two main aspects: species delimitation and diagnosis, based on multiple kinds of data.

The distribution of the tools is also diversified, including command-line tools for those users familiar/comfortable with Python; standalone GUI executables of each module for Windows, Linux, and Mac operating systems for those looking for a single functionality, e.g. a converter, to be called from a single and easily portable file – these tools will necessarily be ‘heavier’ and slightly slower than command-line executables; and a single software package containing all libraries (currently developed for Windows and Linux), from which each module can be launched (Fig. 2). In the future, the latter software package will also enable data transfer between different modules. The GUI software versions are designed to be stable over many different versions of the respective operating systems, e.g., from Windows 7 to Windows 10.

**FIGURE 2.**
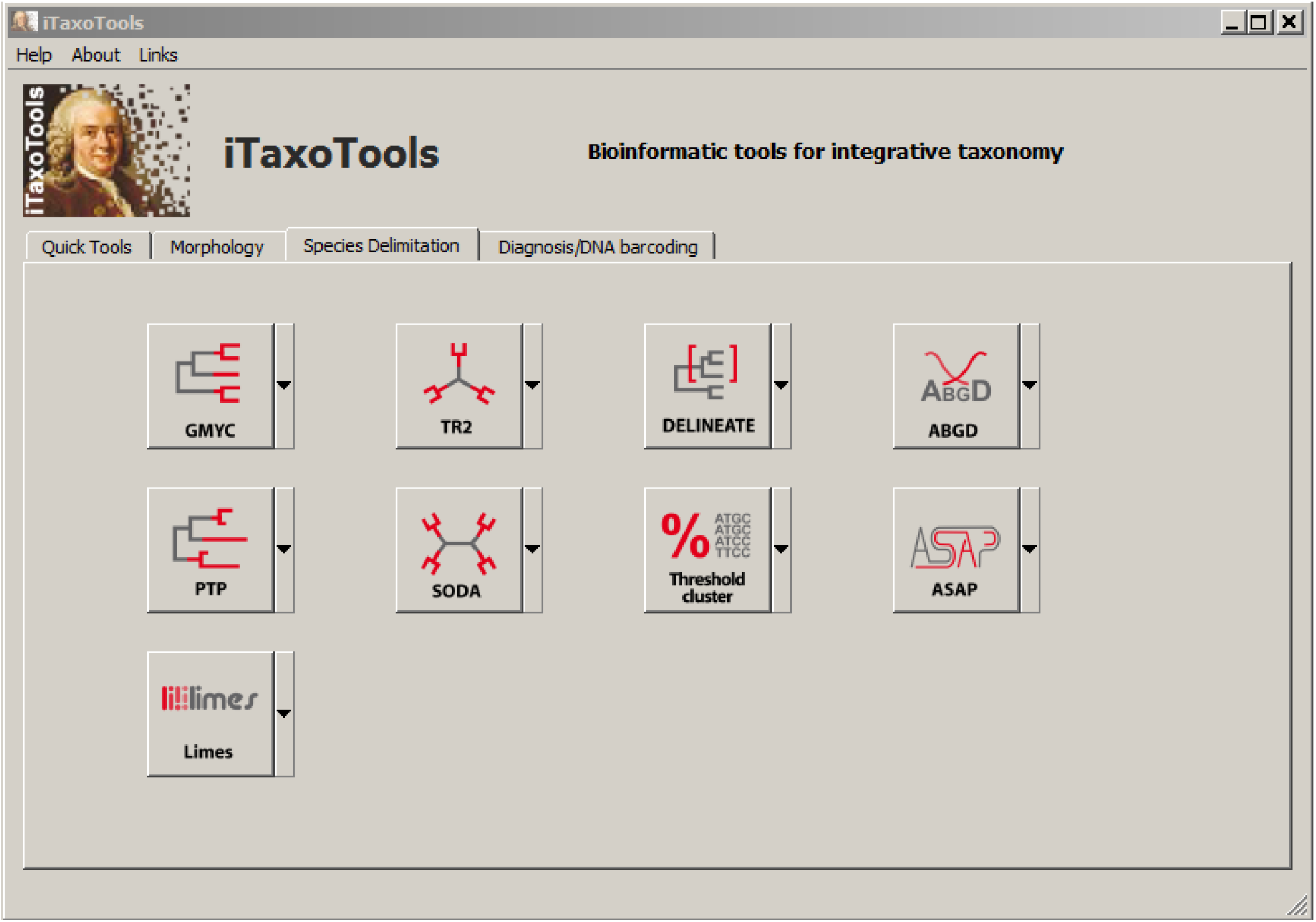
Main launcher window of iTaxoTools 0.1 with the option to start various species delimitation tools (additional tools can be started from the other tabs).

Alpha taxonomy is a primarily **specimen-centered** research field in which specimens – mostly single individual organisms or parts thereof, or cultured isolates composed of multiple individuals – are grouped into species. Consequently, iTaxoTools has implemented tab-delimited text as standard format for most tools, with one column indicating the specimen identifier. This will in subsequent versions allow the user to save the output of different tools for each specimen, and combine these results for further analysis. The tab-delimited format also allows easy editing of the data tables in spreadsheet editors. This specimen-based architecture needed for alpha-taxonomic programs remains valid whether specimens are represented by physical vouchers, images, or in the future maybe by full genome sequences.

Simplicity and **user-friendliness** are at the core of the toolkit we are developing. Because the majority of taxonomists is not familiar with programming languages, such as Python, all our tools are accessible via graphical user interfaces (GUI) – analyses can therefore be carried out with a few intuitive mouse clicks, under default or custom settings, without the need to enter commands in a command line. We also have added autocorrect routines to avoid the loss of time associated with the search for small misspellings or incorrect characters in input files that cause programs to fail. Furthermore, we will provide detailed manuals and wikis with screenshots, along with example files. We chose Python as the main programming language for our package, because it is characterized by its good readability and simple-to-learn syntax, and we documented newly written code extensively, to allow its re-use by other programmers. This comes at the cost of speed that would have been achieved by using the C programming language, but our toolkit in this early phase is not designed to cope with huge genomic datasets or analyses with tens of thousands of specimens. Currently iTaxoTools is designed to provide support for the most common taxonomic research projects that discover and name a limited number of species only (Miralles et al. 2020), but will be extended to large-scale projects in the future.

Considering that powerful programs exist for phylogenetic, phylogenomic and DNA metabarcoding analyses, we did not attempt to include such functionalities in our toolkit. Similarly, we also did not focus on dedicated multiple sequence alignment programs or genome assemblers because (i) these bioinformatic tasks are more efficiently carried out by programs written in C language, (ii) GUI-driven programs and pipelines already exist for alignment and phylogeny (e.g., PAUP, MEGA, PAUP, BEAST: Swofford 2003; Kumar et al. 2018; Bouckaert et al. 2019), genomics, and DNA metabarcoding (e.g., Anslan et al. 2017) and (iii) there is an active community both of commercial companies and academic research teams constantly extending these kinds of programs. We are, however, adding graphical user interfaces and new functionalities to other existing tools that are important for analyses in the context of systematics and that are not yet optimized with user-friendly GUIs. For instance, we have updated the code of Partitionfinder (Lanfear et al. 2016) from Python v. 2 to v. 3, and aim to add a GUI also to the sequence alignment program MAFFT (Katoh & Standley 2013). These developments will be added successively to iTaxoTools.

## Functionalities implemented in iTaxoTools 0.1

Our work on iTaxoTools is ongoing and will be intensified in the period 2021–2023 thanks to support by the DFG SPP 1991 TaxonOmics priority funding program. The current version, published along with the present article, already includes a series of functional tools that we predict will be useful in different steps of the alpha-taxonomic workflow, Data Preparation (mainly Conversion), Analysis, Delimitation, and Diagnosis.

### Data Preparation

Several tools convert among data formats, with the major modules being **dnaconvert** for converting among common DNA sequence formats, **latlonconverter** for converting among geographic coordinate systems (elaborated upon below), and **pyr8s** for converting non-ultrametric trees to ultrametric. A collection of simpler tools includes **fastmerge** and **fastsplit** for splitting and merging large fasta and fastq files, including advanced filtering options by sequence name or sequence motif; **specimentablemerger** and **specimentablepruner** for splitting and merging tab-delimited text files by specimen identifiers; **linebreaker** for converting among Linux and Windows line-break styles (often necessary when processing input files from other bioinformatic tools); **nodenamecorrector** for replacing all non-standard ASCII characters from Newick-format trees; and **unitconverter** for distance, time, volume, molarity, and other units.

**dnaconvert** is a versatile tool to convert DNA (and protein) sequence data among commonly used formats such as fasta, fastq, phylip, or nexus (Fig. 3). Compared to other sequence format converters, dnaconvert is particularly user-friendly in that it autocorrects numerous issues that usually create compatibility problems, e.g., by automatically replacing non-standard ASCII characters from sequence names or auto-renaming sequences in formats of limited sequence name length such as phylip. A main novelty is the support for tab-delimited files because in our experience, it is useful, for small to medium-sized taxonomy projects, to store and organize specimen-based DNA sequence information (DNA barcodes) in spreadsheet editors such as Microsoft Excel or its freeware equivalents Libre Office / Open Office Calc. From these spreadsheets, it is then easy to copy-paste the sequence, specimen-voucher, species and locality columns into dnaconvert and obtain a sequence file for analysis, e.g. in fasta format, with all respective information concatenated in the sequence name. The program also supports a format in which these metadata are bracketed as required for uploading the sequence data along with metadata to the NCBI Genbank repository (i.e., via Submission Portal or BankIt). Lastly, the program also converts Genbank flatfiles into a tabular format, allowing the user to immediately have all relevant metadata associated with the Genbank record in separate columns in a spreadsheet.

**FIGURE 3.**
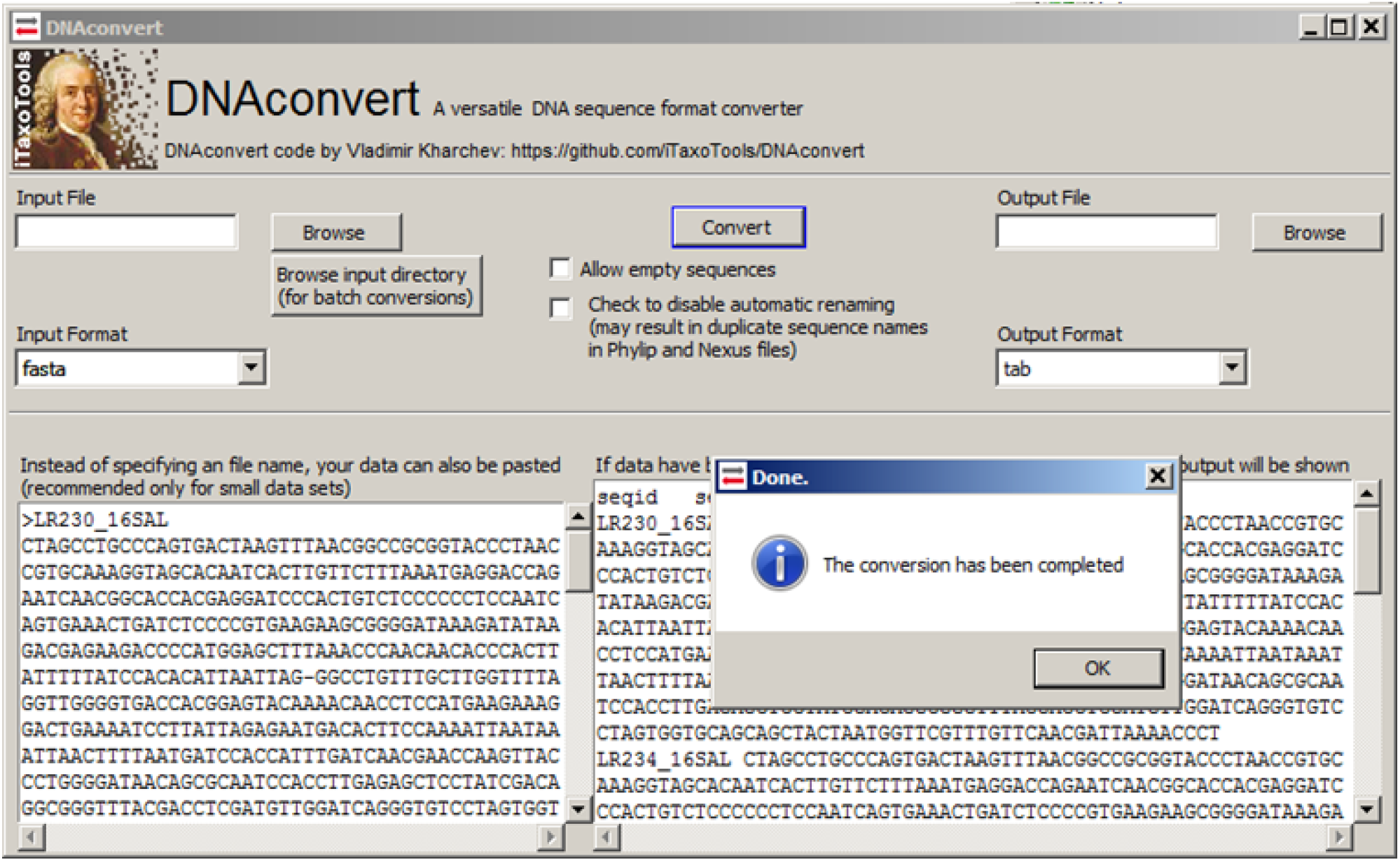
Screenshot of one of the newly programmed quick conversion tools, DNAconvert, which implements numerous autocorrect options to avoid sequence output files generating errors in downstream programs. DNAconvert also supports tab-delimited table input and its conversion to common sequence formats such as FASTA, NEXUS, or PHYLIP, to facilitate storage and management of sequences and sequence metadata in spreadsheet editors such as Microsoft Excel.

**latlonconverter** allows batch conversion of geographic coordinates from a large number of different formats into standard decimal format as required by most geographical information system (GIS) programs. By performing a series of autocorrections of possible typos and then using a heuristic approach, latlonconverter is able to recognize and transform many idiosyncratic formats of geographical coordinates as they are commonly found in specimen databases containing geographical information taken by different researcher. With **spartmapper**, geographical coordinates in combination with a species partition file (spart; Miralles et al. submitted) can be previewed on a map, and then transformed in a kml file that plots all localities on Google Earth and visualizes the geographical distribution of respective species hypothesis (Fig. 4).

**FIGURE 4.**
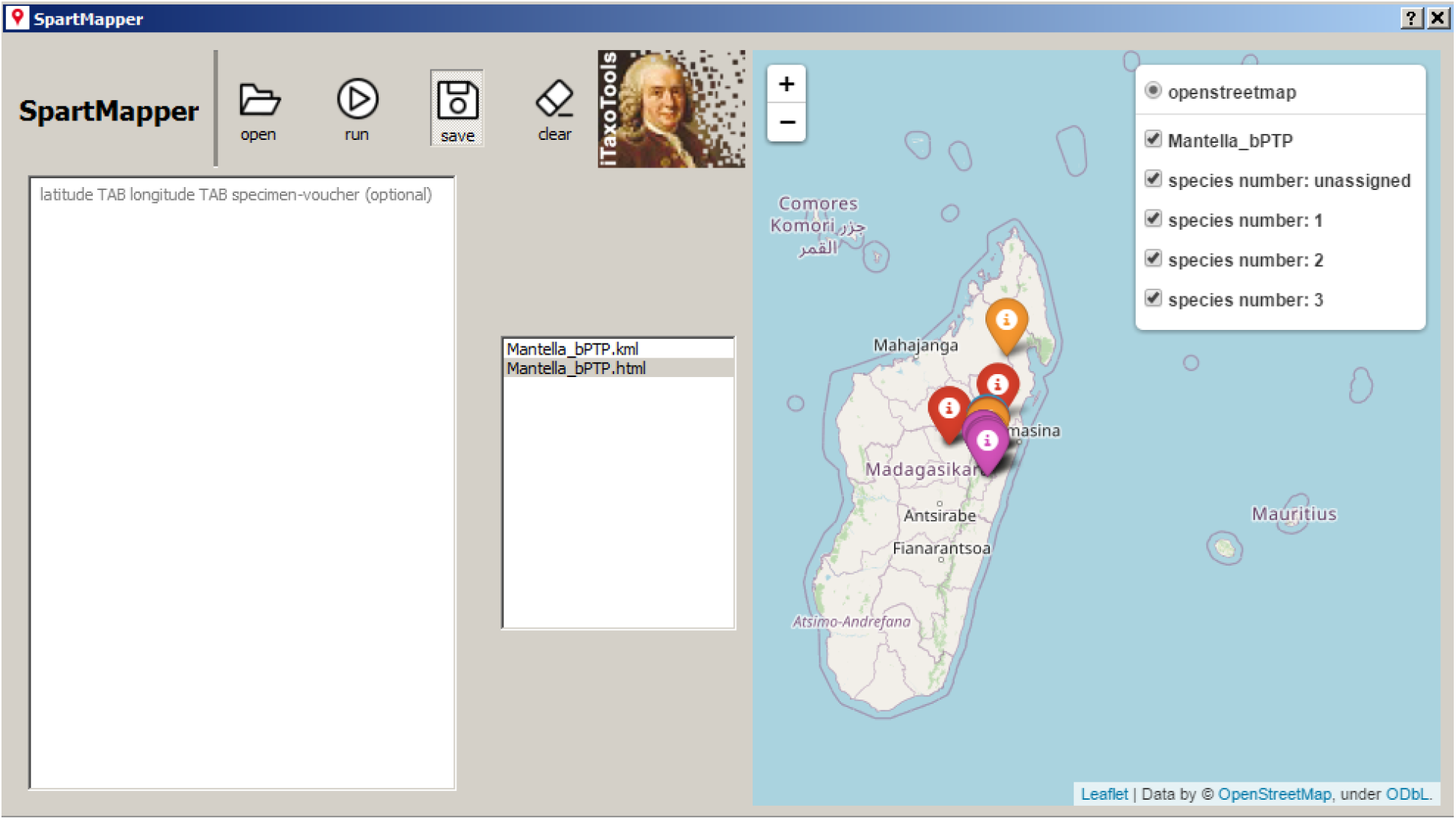
Screenshot of spartmapper, a tool that plots distribution records from geographical coordinates on a map and categorizes the records based on a species hypothesis provided as SPART file (Miralles et al. 2021). The program allows live view and produces a kml file to visualize the records in Google Earth or Google Maps.

**pyr8s** is one of our flagship modules (Fig. 5). For many evolutionary analyses, but also for species delimitation (e.g. GMYC), ultrametric phylogenies are required where non-ultrametric trees are available. This conversion is rather complex and can be time-intensive. While numerous programs exist to calculate time trees (e.g., MCMCtree, BEAST, MEGAX: Yang & Rannala 2006; Bouckaert et al. 2019; Kumar et al. 2018), they usually require DNA sequence information in addition to a previously inferred phylogenetic tree. For iTaxoTools, we opted to recuperate a vintage approach, non-parametric rate smoothing (NPRS), initially developed by Sanderson (1997) and later implemented as part of the program r8s (Sanderson 2003). This method only requires a phylogenetic tree as input, with the option to add one or more time calibration points. NPRS has previously been implemented in the R package ape (Paradis et al. 2004), but was removed from the latter and from the newest releases of r8s due to licensing issues. Specifically, the original version of r8s relied on a modified implementation of Powell’s conjugate direction method which was incompatible with open-source licensing (Powell 1964; Gill et al. 1981; Press et al. 1992). In the GUI-driven tool **pyr8s**, the NPRS algorithm has been newly coded, making use of the open-source libraries DendroPy (Sukumaran & Holder 2010) and SciPy (Virtanen et al. 2020), thus resolving the previous licensing issues. This new version provides a GUI for user-friendly setting of time constraints, exposes a Python interface for lower-level analysis and maintains support for r8s-formatted input files.

**FIGURE 5.**
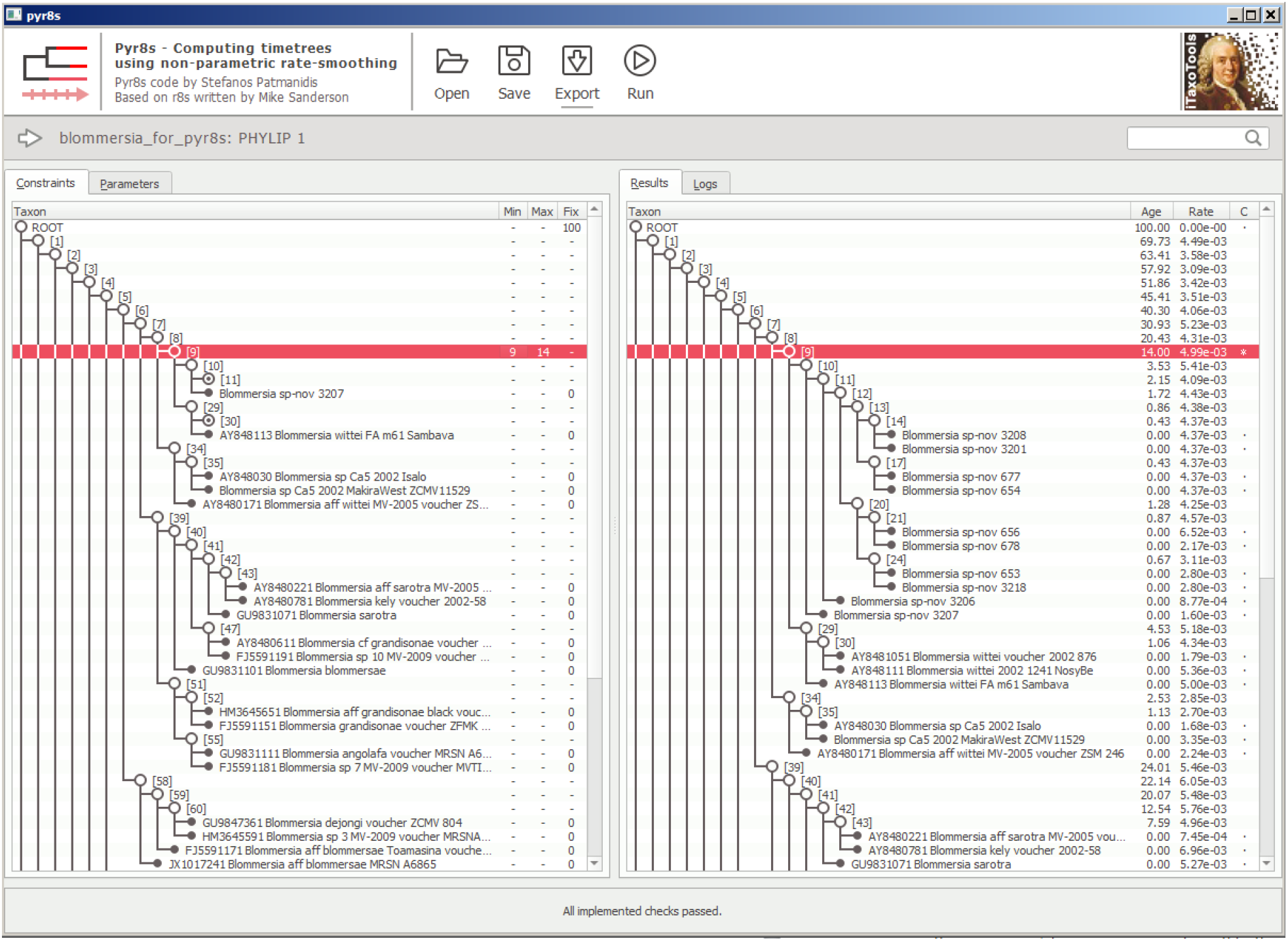
Screenshot of pyr8s after running an analysis and converting a tree into an ultrametric timetree. The red-highlighted row marks a node for which age constraints had been set before the analysis.

### Analysis

We include several data analysis modules: **TaxI2** for calculation of pairwise distances among individuals, and **morphometricanalyzer** for basic morphometric analyses (elaborated upon below). For convenience, we also include **simplestatscalculator**, a utility for quick, basic statistical analyses of manually entered or pasted data.

**TaxI2** is a tool for pairwise sequence comparison. To analyze DNA barcoding data sets, Steinke et al. (2005) proposed the program TaxI, which performs pairwise alignments between sequences and calculates pairwise distances based on these alignments. Compared to a multiple sequence alignment (MSA) the authors argued that these distance calculations may be more accurate in the case of highly divergent markers including multiple insertions and deletions, such as stretches of mitochondrial ribosomal RNA genes. The pure-Python tool **TaxI2** performs similar calculations, with numerous added functionalities such as support for pre-MSA aligned data sets. The tool has two main analysis modes: First, following the original TaxI approach, it can compare a set of sequences against a reference database, via pairwise alignments, identifies for each query the closest reference sequence, and calculates various genetic distances among the two. Second, it also can perform all-against all comparisons of a set of sequences. In this latter approach, sequences can be added in tab-delimited table format along with species name, and from these data the program calculates within-species, between-species, and between-genus distances. Various metrics and graphs defining the barcode gap in a given data set are also included in the output. The program furthermore performs a simple threshold-based clustering of DNA sequences into OTUs, following the approach previously implemented in TaxonDNA (http://taxondna.sourceforge.net/; Meier et al. 2006), and outputs the resulting species partition as SPART file (Miralles et al. 2021)

**morphometricanalyzer** is our tool for exploratory analysis of morphometric datasets. Integrative taxonomists do not only use molecular data. In many cases, a limited number of one-dimensional morphometric measurements such as body length and width (or leaf length and width in plants) are taken and compared among groups of individuals. For simple statistical analyses, we have included the tool **morphometricanalyzer** which performs a series of exploratory routine comparisons from morphometric data. It takes as input tab-delimited text files with species hypotheses and a series of other optional categories, and then performs automatically a series of statistical comparisons between species (and between other categories), such as calculations of means, medians, standard deviation, minimum and maximum values; pairwise Mann-Whitney U-tests and Student’s t-tests between all pairs of species; a simple Principal Component analysis; and calculation of ratios among original values as a means to size-correct them, followed by statistical comparison of these size-corrected values. Finally, the program also outputs pre-formulated taxonomic diagnoses, with full-text sentences specifying by which morphometric value or ratio a species/population differs from other species/populations with statistical significance, or without value overlap. It would also be possible to explore non-morphological (e.g. bioacoustic) data with this tool, although it is primarily developed for morphometrics.

### Delimitation

A special emphasis in the first development phase of iTaxoTools is species delimitation, a burgeoning field in systematics. The available species delimitation algorithms mostly use DNA sequence data and tend to overestimate the number of species in a data set (e.g., Miralles et al. 2013); indeed, they may delimit populations rather than species (Sukumaran & Knowles 2017). Yet, such automated delimitation may play a role in formulating initial species hypotheses that can then be tested in an integrative taxonomy pipeline. In the first version of iTaxoTools, we have focused on tools already available in Python programming language. For these tools, we added user-friendly GUIs and slightly extended the functionality, for example by enabling them to output species partition information in the standardized “spart” format proposed by Miralles et al. (2021). The current version of iTaxoTools includes GUI-enhanced versions of **PTP** (Zhang et al. 2013) (Fig. 6) and **GMYC** (Pons et al. 2006; Fujisawa & Barraclough 2013; Python version J. Zhang) which delimit species from single-locus trees; **tr2** (Fujisawa et al. 2016) and **DELINEATE** (Sukumaran et al. 2020) that use coalescence-based approaches on multiple gene trees; and **ABGD** (Puillandre et al. 2012) (Fig. 7) and **ASAP** (Puillandre et al. 2020) that are alignment-based and rely on calculations of genetic distances. For some of these tools (PTP, GMYC, tr2, DELINEATE) the current pre-release GUI versions are still basic and only run under default settings; options to change and refine parameters will be added to the first complete release. iTaxoTools also includes **LIMES 2.0**, a program to handle and compare species partitions obtained by these various approaches (Ducasse et al. 2020, Miralles et al. 2021).

**FIGURE 6.**
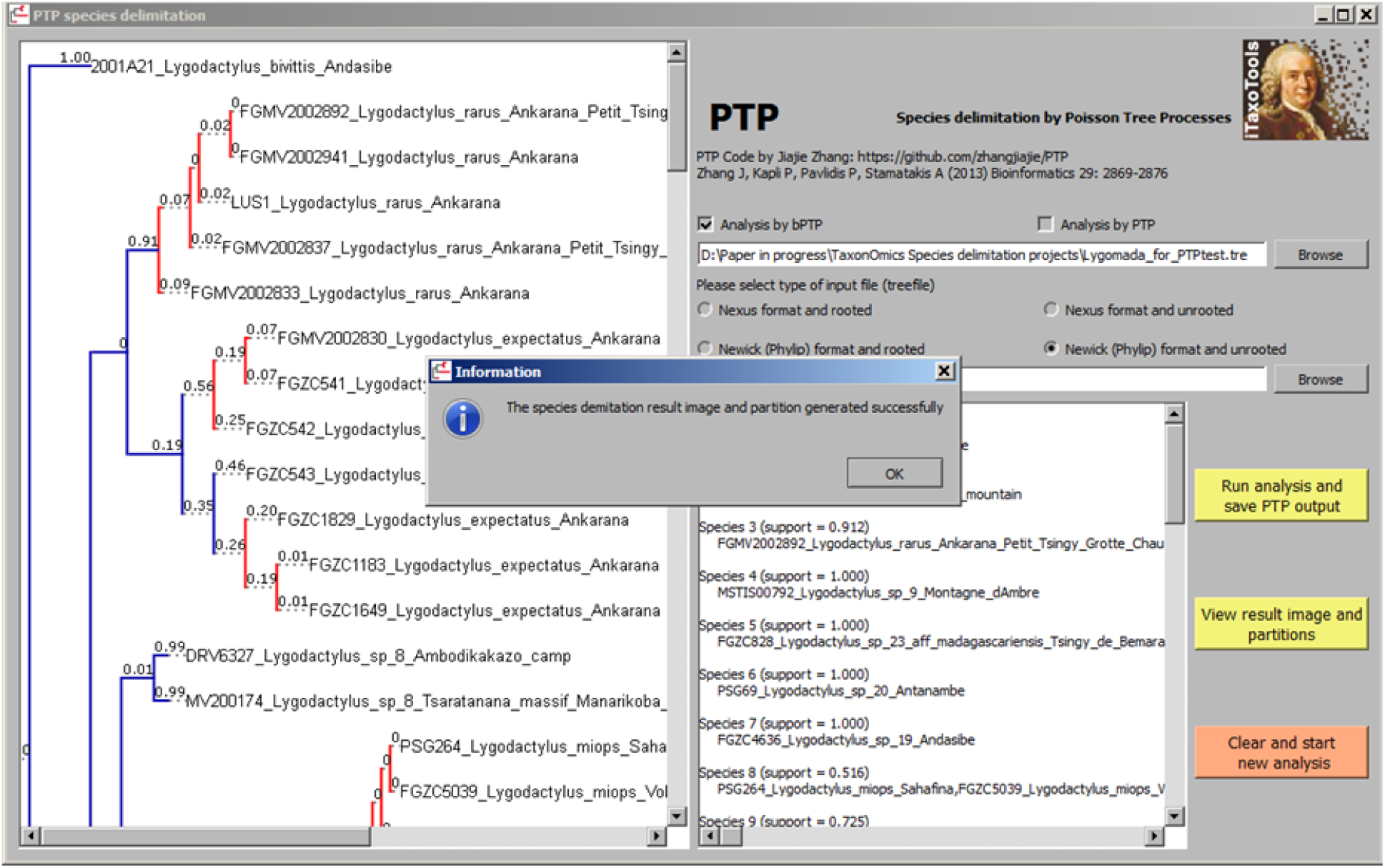
Screenshot of the GUI-based version of PTP, a program that delimits species from non-ultrametric trees. The original Python code of PTP was written by Zhang et al. (2013); iTaxoTools adds the GUI, as well as functionality to export species partition in the SPART format (Miralles et al. 2021).

**FIGURE 7.**
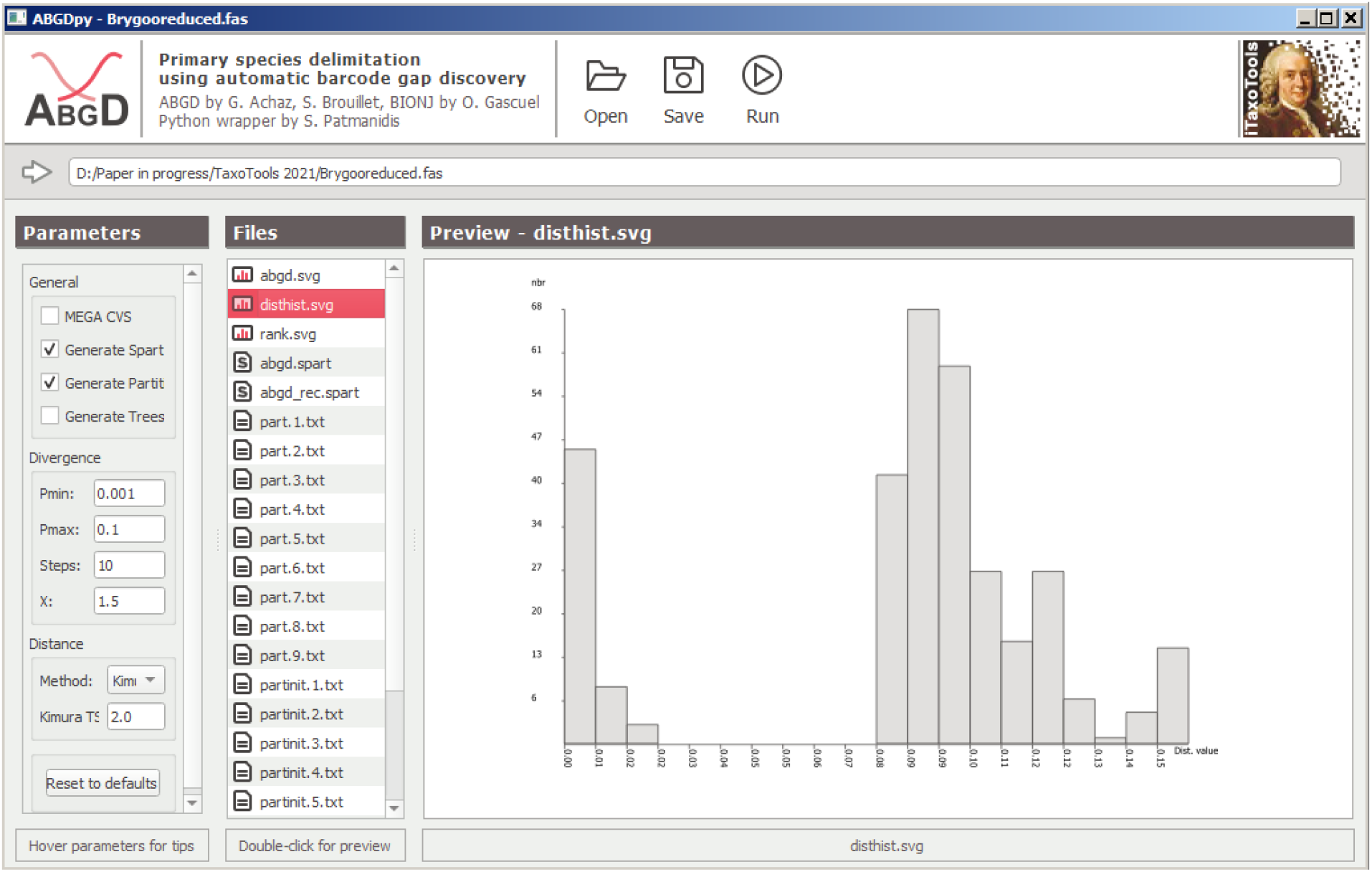
Screenshot of the GUI-based version of ABGD, a program that delimits species by detecting the barcoding gap from pairwise single-locus sequence distances (Puillandre et al. 2012). For this tool, the original ABGD code written in C was wrapped with a Python GUI and compiled as standalone executable. The different output files produced by ABGD (text and graphs) can be selected and pre-viewed within the GUI.

### Diagnosis

The diagnosis of new species – rather than its lengthy description – represents the most important part of the alpha-taxonomic process, and in all Nomenclatural Codes, diagnosis can be based on molecular, as well as morphological characters (Renner, 2016). Several software tools have been proposed to extract diagnostic nucleotide positions of clades and species, either phylogeny-based (caos; Sarkar et al. 2008) or primarily alignment-based (MolD, Fastachar, DeSignate: Fedosov et al. 2019; Merckelbach & Borges 2020; Hütter et al. 2020). In order to facilitate the use of such DNA characters in differential diagnoses of new species, we implemented a crucial new tool for DNA taxonomy named **dnadiagnoser**. Compared to other tools, dnadiagnoser has various functionalities to improve the use of DNA characters in species diagnosis. It takes as input tab-delimited text files in which one column specifies the unit for analysis (typically the species), and provides as output pre-formulated text sentences which specify (i) in a pairwise fashion, all the diagnostic sites of one species against all other species, and (ii) the unique diagnostic sites (if any) that differentiate a species against all other species. These text sentences can then directly be used in species diagnoses. As a further innovation dnadiagnoser interprets one of the sequences in the input alignment as reference sequence and outputs the diagnostic sites relative to this sequence. To facilitate such comparisons, the program also includes a series of standard reference sequences (such as the full *Homo sapiens* COI or *cox1* gene) and allows as input unaligned sequences, which are then pairwise aligned against the reference sequence to identify diagnostic positions and label them according to their position in the reference sequence, a procedure that works reliably in sets of sequences with no or only few insertions or deletions such as COI. In addition, we have also programmed a GUI for **MolD** (Fedosov et al. 2019), a program that is tailored for recovering DNA-based diagnoses in large DNA dataset, and is capable of identifying diagnostic combinations of nucleotides (DNCs) in addition to single (pure) diagnostic sites. The crucial and unique functionality of MolD allows assembling DNA diagnoses that fulfil pre-defined criteria of reliability, which is achieved by repeatedly scoring diagnostic nucleotide combinations against datasets of in-silico mutated sequences.

## Future extensions

Our goal with this paper is to make the tools we have developed available to the community as soon as possible so they may be critically evaluated and improved. The next developments will be in three fields: (i) **Geography:** iTaxoTools will not compete with geographical information systems (GIS), but there are a number of recurrent and rather simple geographical analyses in alpha-taxonomy that can be facilitated by bioinformatic tools, in particular calculation of linear distances among sites and of the surface (minimum convex polygon) of a distribution range of a species, based on a set of georeferenced locality points, and most importantly, a simple graphical editor that outputs publication-ready distribution maps, with customizable colors and symbols for different species, from a set of georeferenced locality records. For more sophisticated analyses, connecting iTaxoTools (via data formats such as SPART) with dedicated toolboxes for analysis of spatial biodiversity data such as SDMToolbox (Brown et al. 2017) could allow e.g. for comparative niche modelling of alternative species partitions. (ii) **Extraction of diagnostic traits from specimen data:** Besides molecular diagnosis with MolD and dnadiagnoser, we plan to develop a tool that automatically outputs diagnoses based on (specimen-based) categorical data sets of morphological characters. (iii) **Connection to other programs:** We also plan to explore options to connect iTaxoTools to the DELTA (DEscription Language for TAxonomy) software package (Coleman et al. 2010). DELTA is a format for coding descriptive taxonomic information that however is primarily species-based (not specimen-based as iTaxoTools), and a series of programs have been developed on this basis, spearheaded by M. Dallwitz at CSIRO (Canberra, Australia) (Dallwitz 1974, 1980). The new Free DELTA platform launched in 2000 (http://freedelta.sourceforge.net/) includes options for editing and maintenance of data sets in DELTA formats, as well as utilities for data conversion, interactive identification of taxa, automated generation of diagnostic keys, and descriptions. Especially, information on species-specific molecular and morphological characters identified in iTaxoTools could be seamlessly coded in DELTA, making use of pydelta (http://freedelta.sourceforge.net/pydelta/).

The biggest gap in taxonomy software so far is the integrative aspect in the sense of Dayrat’s (2005) concept of integrative taxonomy (see also Padial et al. 2010). That is, the many available species delimitation programs all output a species hypothesis based on one analytical approach – usually based on only a molecular data set, with a few exceptions such as iBPP, which can integrate morphometric and molecular data (Solís-Lemus et al. 2015). Approaches that combine information from different lines of evidence into species delimitation are exceedingly scarce. As an example, DELINEATE (Sukumaran et al. 2020) allows the user to fix a series of species hypotheses (i.e., firmly assign a series of specimens to species) while letting the other specimens “float” freely in the analysis and assign them to either one of the previously defined species, or to a new species. Such an option of “prior” species delimitation should be universally available to users as a manual option (e.g., if evidence comes from field or experimental data on hybridization, genomic information, or other data that are yet difficult to code or implement in species delimitation software), similar to what is implemented in DELINEATE. But ideally, automated proposals of firm *a priori* evidence for two specimens to either belong to two species, or to the same species, could also be elaborated by the software – for instance, using evidence such as sympatric geographical occurrence without gene flow, full concordance between genetic and morphological characters, or exceedingly high genetic distances. We plan to develop concepts for such analysis priors, and start implementing them in a iTaxoTools webserver pipeline, in the next years.

Importantly, our project is open for other developers to join, and for the taxonomic community as a whole to provide suggestions. We especially welcome proposals of additional tools that could help to streamline and accelerate the whole process of delimiting and naming species (whether it concerns the initial step of data acquisition, their treatment, their analyses, or their final submission to a dedicated repository). Only practicing taxonomists know which parts of the alpha-taxonomic workflow for their group of taxa is particularly time-consuming, and where time and effort is lost with repetitive, manual tasks that could be as well automatically performed by a computer program – and thereby formulate requirements for such dedicated programs.

## Perspectives for iTaxoTools

The different taxonomic tools made available here are performing analyses offline on a local computer (and in the future will also be available on a webserver), but without linking to external resources. True next-generation taxonomy will require linking specimen-based taxonomy software with online resources and databases, and scaling the analyses to data of many thousands of specimens. On the one hand, this involves archiving newly acquired data in dedicated repositories (Miralles et al. 2020). But on the other hand, it means aggregating for each specimen identifier DNA sequences, morphological characters, images, and increasingly -omics data (e.g., Lendemer et al. 2020), and then entering these large-scale cyberspecimen data into species delimitation, diagnosis and naming pipelines. The process could be coupled with machine-learning programs to automatically extract diagnostic traits e.g. from images, with data aggregators such as GBIF (gbif.org) and online tools such as Map of Life (mol.org), or Timetree of Life (timetree.org) to obtain geographical and temporal context, and distribution models for alternative species hypotheses. These bioinformatic opportunities may gain power under a view of species as probabilistic hypotheses that may allow defining probability thresholds of integrative taxonomic analysis above which lineages can be confidently named as species by semi-automated pipelines. While the current version of iTaxoTools is far from this vision, it may represent a seed for developing the necessary environment, and a sandbox to test software tools with the potential to significantly accelerate the inventory of life.

## Acknowledgments

We are grateful to Mike Sanderson for information on the initial code of r8s. This study was supported by the Deutsche Forschungsgemeinschaft (grant VE247/16-1 – HO 3492/6-1 and RE 603/29-1) in the framework of the ‘TaxonOmics’ priority program.

## Notes

### Competing Interest Statement

The authors have declared no competing interest.

